# Characterization of Brain-Derived Extracellular Vesicle Lipids in Alzheimer’s Disease

**DOI:** 10.1101/2020.08.20.260356

**Authors:** Huaqi Su, Yepy H. Rustam, Colin L. Masters, E Makalic, Catriona McLean, Andrew F. Hill, Kevin J. Barnham, Gavin E. Reid, Laura J. Vella

## Abstract

Lipid dyshomeostasis is associated with the most common form of dementia, Alzheimer’s disease (AD). Substantial progress has been made in identifying positron emission tomography (PET) and cerebrospinal fluid (CSF) biomarkers for AD, but they have limited use as front-line, non-invasive diagnostic tools.

Small extracellular vesicles (EVs) are released by all cell types and contain an enriched subset of their parental cell molecular composition, including lipids. EVs are released from the brain into the periphery, providing a potential source of tissue and disease specific lipid biomarkers. However, the EV lipidome of the central nervous system (CNS) is currently unknown and the potential of brain-derived EVs (BDEVs) to inform on lipid dyshomeostasis in AD remains unclear. The aim of this study was to reveal the lipid composition of BDEVs in human frontal cortex tissue, and to determine whether BDEVs in AD have altered lipid profiles compared to age-matched neurological controls (NC).

Here, using semi-quantitative mass spectrometry, we describe the BDEV lipidome, covering 4 lipid categories, 17 lipid classes and 692 lipid molecules. Frontal cortex-derived BDEVs were enriched in glycerophosphoserine (PS) lipids, a characteristic of small EVs. Here we report that BDEVs are enriched in ether-containing PS lipids. A novel finding that further establishes ether lipids as a feature of EVs.

While no significant changes were detected in the frontal cortex in AD, the lipid profile of the BDEVs from this tissue exhibited disease related differences. AD BDEVs had altered glycerophospholipid (GP) and sphingolipid (SP) levels, specifically increased plasmalogen glycerophosphoethanolamine (PE-P) and decreased polyunsaturated fatty acyl containing lipids (PUFAs), and altered amide-linked acyl chain content in sphingomyelin (SM) and ceramide (Cer) lipids relative to vesicles from neurological control subjects. The most prominent alteration being a two-fold decrease in lipid species containing docosahexaenoic acid (DHA).

The in-depth lipidome analysis provided in this study highlights the advantage of EVs over more complex tissues for improved detection of dysregulated lipids that may serve as potential biomarkers in the periphery.

## Introduction

Alzheimer’s disease (AD) is a neurodegenerative disorder and the most common form of dementia. AD has an extended preclinical window which offers hope in treating the disease, however clinical differentiation and diagnosis remains difficult in the absence of blood-based biomarkers and current therapies are unable to halt disease progression.

Lipids are involved in maintenance of cellular homeostasis and are key components of cellular membranes. They have multiple functional roles, including regulation of energy storage and signal transduction, and also for the segregation of chemical reactions in discrete organelles. It has previously been reported that lipid metabolism is extensively reprogrammed in AD as detected in brain, cerebrospinal fluid (CSF) and blood (Han et al., 2011, He et al., 2010, Kosicek and Hecimovic, 2013, Mielke et al., 2014, Mielke and Lyketsos, 2010, Wong et al., 2017, Wood, 2012, Mielke et al., 2010a, Mielke et al., 2010b, Mielke et al., 2011). Glycerophospholipids (GP), as the most abundant structural components within cellular membranes, regulate membrane mobility, provide secondary messengers for cellular signalling, play a critical role in synaptic transmission between neurons and can cause neuronal death when dysregulated (Bennett et al., 2013, Wong et al., 2017). The homeostasis of GP has been reported to be altered in AD, with a general downregulation of glycerophosphocholine (PC), glycerophosphoethanolamine (PE) and glycerophosphoinositol (PI) species (Bennett et al., 2013, Kosicek and Hecimovic, 2013, Wong et al., 2017, Wood, 2012), with additional alterations in a specific GP subclass, namely alkenyl (i.e., plasmalogen)-containing species in AD brain (Ginsberg et al., 1995, Han et al., 2001, Pettegrew et al., 2001). Sphingolipids (SP), as another fundamental structural component of cellular membranes, and comprised of sphingomyelin (SM), ceramide (Cer) and other glycosphingolipid species, play a critical role in the central nervous system (CNS) and are heavily involved in neuronal signalling (Wood, 2012). SM and Cer are major components of the myelin sheath, and deregulation of the SM/Cer signalling cascade causes synaptic dysfunction in AD, while up-regulation of Cer results in neuro-inflammation and neuronal apoptosis (Cutler et al., 2004, Haughey et al., 2010, He et al., 2010, Mielke and Lyketsos, 2010, Satoi et al., 2005, Wong et al., 2017, Wood, 2012). Moreover, decreases in SM and increases in Cer are accompanied by the over-expression of two lipid regulating enzymes in the SP pathway, namely acid sphingomyelinase and acid ceramidase, in human AD brain (He et al., 2010). However, the relative levels of lipids reported in previous studies are partially conflicting, likely due to the differences in brain region, disease severity and comorbidity, etc. (Han et al., 2002, Bandaru et al., 2009, Cutler et al., 2004, Han et al., 2001, He et al., 2010, Kosicek and Hecimovic, 2013, Pettegrew et al., 2001, Wong et al., 2017, Wood, 2012).

Exosomes are small extracellular vesicles (EVs) released from cells upon fusion of the multi-vesicular body (MVBs) with the plasma membrane (Johnstone et al., 1987, Raposo et al., 1996). They are surrounded by a lipid bilayer, and packaged with cargo including proteins, lipids, nucleic acids and metabolites which reflect their cellular origin and mirror the physiological or pathological condition of the parental cell (Kanninen et al., 2016). EVs have a unique lipid composition with multiple studies reporting enrichment of specific lipid classes including SM, glycerophosphoserine (PS), and PC and PE ether lipids, in EVs relative to their parental origin (Hessvik and Llorente, 2017, Kanninen et al., 2016, Laulagnier et al., 2004, Lydic et al., 2015, Record et al., 2014, Record et al., 2018, Llorente et al., 2013, Phuyal et al., 2015, Simbari et al., 2016). EVs are regarded as critical players in intercellular communication by inducing phenotypic changes and altering homeostasis in recipient cells upon fusion or uptake (Rai et al., 2019, Record et al., 2014, Simons and Raposo, 2009). Given EVs are present in the extracellular environment, including bodily fluids, there is intense interest in using EVs as a source of tissue and disease specific biomarkers. Recent studies have revealed exosomal proteins as potential biomarkers in non-small cell lung cancer and pancreatic cancer (Li et al., 2017, Melo et al., 2015) and certain lipid species in urinary exosomes have been reported to serve as biomarkers for prostate cancer diagnosis (Skotland et al., 2017a). While the proteomic and nucleic acid compositions of EVs have been extensively studied, lipidomic profiling of EVs is still in its infancy (Skotland et al., 2020) even though lipids are an essential component of EVs.

As BDEVs are predicted to participate in AD pathogenesis (DeLeo and Ikezu, 2018, Kanninen et al., 2016, Rajendran et al., 2006, Sharples et al., 2008, Thompson et al., 2016, Vella et al., 2016, Vingtdeux et al., 2012), we hypothesized that EVs could provide a rich source of lipids that reflect alterations observed in the CNS in AD (He et al., 2010, Wood, 2012). Here, using methods we previously developed for the isolation and enrichment of EVs from human brain tissue (Vella et al., 2017), coupled with semi-quantitative ultrahigh resolution accurate mass spectrometry (UHRAMS) based lipidome analysis (Ryan and Reid, 2016), we have determined, for the first time, the lipid composition of BDEVs in human frontal cortex and identified the lipidomic signature of EVs in human AD compared to gender and age-matched neurological controls (NC).

## Materials and Methods

### Human frontal cortices

Fresh frozen human post-mortem frontal cortex tissues of n=8 AD male subjects (mean age 74.5 ± 7.0 years) and n=8 gender and age-matched NC subjects (mean age 73.5 ± 5.9 years) with no evidence of neurological disease, stored at −80°C, were obtained from the Victoria Brain Bank. The average post mortem delay before tissue collection was 23.3 ± 17.4 hrs for AD and 42 ± 16.3 hrs for NC. Individual tissue case information, including age, PMI and ApoE genotype, is provided in **Supplemental Table 1**. Tissue handling and experimental procedures were approved by The University of Melbourne human ethics committee and in accordance with the National Health and Medical Research Council guidelines.

### BDEV enrichment

The BDEV isolation method was modified from our previously published protocol (Vella et al., 2017). A schematic of the workflow is shown in **Supplemental Figure 1**. Frozen frontal cortex tissues (approximately 2 g) were sliced lengthways on ice to generate 1–2 cm long, 2–3 mm wide tissue sections. Approximately 30 mg tissue pieces from each sample (“Brain Total”) were collected, weighed and placed in 19x volume of tissue weight of Dulbecco’s phosphate buffered saline (DPBS, Thermo Fisher Scientific) solution containing 1x PhosSTOP™ phosphatase inhibitor (Sigma Aldrich) / cOmplete™ protease inhibitor (including EDTA, Sigma Aldrich) for immunoblot analysis. The remaining cut tissue sections were weighed and incubated with 50 U/mL collagenase type 3 (#CLS-3, CAT#LS004182, Worthington) digestion buffer (at ratio of 8μL / mg tissue) in a shaking water bath (25°C, a total of 20 minutes). During incubation, tissue slices were inverted twice at the 10-min time point, gently pipetted up and down twice at the 15-min time point and then allowed incubation for a further 5 minutes, followed by the addition of ice-cold 10x inhibition buffer, which was made of 10x phosphatase inhibitor and 10x protease inhibitor in DPBS. The final concentration of inhibition buffer in solution was 1x. The dissociated tissue in solution was subjected to a series of centrifugations, including a 300 x g, 4°C for 5 min, a 2000 x g, 4°C for 10 min and a 10,000 x g, 4°C for 30 min. Representative 300 x g pellets were collected (“Brain+C” for collagenase treatment) and either placed in 19x volume of tissue weight of DPBS with 1x phosphatase inhibitor / protease inhibitor solution for protein quantification and immunoblot analysis or combined with 19x volume of tissue weight of ice-cold 60% methanol (LCMS grade, EMD Millipore Corporation) containing 0.01% (w/v) butylated hydroxytoluene (BHT, Sigma Aldrich) for lipid extraction. The 10,000 x g supernatant was loaded on top of the triple sucrose density gradient (0.6 M, 1.3 M, 2.5 M) as indicated in the method (Vella et al., 2017) in ultra-clear SW40Ti tubes (Beckman Coulter). The sucrose gradients were centrifuged at 200,000 x g avg at 4°C for 173 min using a SW40Ti rotor (Beckman Coulter). After the spin, the three fractions (F1, F2 and F3, 1.2 mL each) were sequentially collected and refractive index was measured. Each fraction was subjected to a wash spin in ice-cold DPBS at 128,000 x g avg, at 4°C for 80 min using a F37L-8×100 rotor (Thermo Fisher Scientific). The pelleted EVs were resuspended in 150 μL ice-cold DPBS with 1x phosphatase inhibitor / protease inhibitor solution. In each 150 μL vesicle suspension, 5 μL were aliquoted for TEM, 65 μL were aliquoted for protein quantification and immunoblot analysis and 80 μL were used for lipid extraction.

### Homogenization and protein quantification for immunoblot

Tissue in DPBS solution with 1x phosphatase inhibitor / protease inhibitor was homogenised for 12 sec at 50% intensity using a tapered microtip attached to the Sonifier Cell Disruptor (Branson) and sonicated in ice-cold water bath for 20 min before submitted to clarification spin (10,000 x g, 10 min, 4°C). The EV suspensions were sonicated in ice-cold water bath sonicator for 20 min. Protein content in tissue homogenates and EV suspensions were determined by Pierce™ BCA assay kit (Thermo Fisher Scientific) according to the manufacturer’s instructions.

### SDS-PAGE and Western Blot (WB) analysis

Samples were prepared in 4 x Laemmli sample buffer, then boiled (10 minutes, 90°C) followed by centrifugation (14,000 x g, 1 min). Normalised samples (1 – 3 µg) were electrophoresed on 12% Mini-Protean TGX Stain-Free gels (BioRad) or 4-20% Criterion TGX Stain-Free Precast gels (BioRad) in Tris/Glycine/SDS running buffer (BioRad) for 30 minutes at 245 V. Proteins were transferred onto nitrocellulose membranes using either a Trans-Blot® Turbo™ Transfer System (7 minutes, 25V, BioRad) or an iBlot™ 2 Dry Blotting System (P0 method, Thermo Fisher Scientific). Membranes were blocked with 5% (w/v) skim milk in TBS-T (1 hour, room temperature) followed by overnight incubation with primary antibodies (calnexin #ab22595 from Abcam, syntenin #ab133267 from Abcam, TSG101 #T5701 from Sigma) at 4°C. Membranes were washed 4 times with TBS-T (30 min, room temperature) following a 1 hr, room temperature incubation of the secondary anti-rabbit antibody either conjugated to HRP (#7074S from Cell Signalling Technology) or the IRDye® 800CW Goat anti-Rabbit IgG secondary antibody (#925-32211, LICOR). All antibodies were diluted in 5% (w/v) skim milk in TBS-T. Membranes were washed 4 times with TBS-T (30 min, room temperature). The membranes were visualised either on a ChemiDoc MP Imager (BioRad) following development with Clarity™ Western Enhanced Chemiluminescence Blotting Substrate (BioRad) or on an Odyssey® Fc Imaging System (LI-COR).

### Transmission electron microscopy (TEM)

A 5 μL EV suspension from F1, F2 or F3 in 1% (w/v) electron microscopy-grade glutaraldehyde was absorbed onto neutralised 300-mesh carbon-coated formvar copper grids (ProSciTech, QLD, Australia) for 5 min. Excessive liquid was removed and grids were washed with DPBS and MilliQ water, and negatively stained with 2% (w/v) saturated aqueous uranyl acetate for 12 sec. Excessive stain was removed and grids were dried. Images were taken on a Tecnai G2 F30 (FEI, Eindhoven, The Netherlands) transmission electron microscope operating at 300 kV wide field images encompassing multiple vesicles were captured to provide an overview of the fraction in addition to close up images. Electron microscopy was performed in the Bio21 Advanced Microscopy Facility, Bio21 Molecular Science and Biotechnology Institute, at The University of Melbourne.

### Monophasic lipid extraction from tissues and derived exosomes

The “total brain with collagenase” tissue pellets in ice-cold 60% methanol containing 0.01% (w/v) BHT were homogenised using a cell disrupter as described above. 100 μL of the homogenates were combined with 100 μL of 60% methanol containing 0.01% (w/v) BHT. 80 μL of the F2 BDEV suspensions were combined with 20 μL of ice cold methanol with 0.1% (w/v) BHT and 100 μL of ice-cold methanol to make a final volume of 200 μL 60% methanol containing 0.01% (w/v) BHT. All samples were sonicated in an ice cold water bath sonicator (20 min) prior to lipid extraction. Monophasic lipid extraction followed the method previously reported by Lydic *et al*. (Lydic et al., 2015) with modification as described below. 120 μL of MilliQ water, 420 μL of methanol with 0.01% (w/v) BHT, and 270 μL of chloroform were added to all samples. For every 10 μg protein present in the samples, 1 μL of a customised isotope labelled internal standard lipid mixture and 1 μL of a d5-TG Internal Standard Mixture I (Avanti Polar Lipids, Alabaster, AL, USA) were added. The customised isotope labelled internal standard mixture was comprised of 14 deuterated lipid standards (Avanti Polar Lipids, Alabaster, AL, USA): 15:0-18:1(d7) PC (250 µM), 15:0-18:1(d7) PE (240 µM), 15:0-18:1(d7) PS (250 µM), 15:0-18:1(d7) PG (20 µM), 15:0-18:1(d7) PI (220 µM), 15:0-18:1(d7) PA (180 µM), 18:1(d7) LPC (45 µM), 18:1(d7) LPE (10 µM), 18:1(d7) Chol Ester (10 µM), 18:1(d7) MG (10 µM), 15:0-18:1(d7) DG (17 µM), 18:1(d9) SM (80 µM), d18:1(d7)-15:0 Cer (40 µM) and Cholesterol(d7) (20 µM). The d5-TG Internal Standard Mixture I contained 20:5-22:6-20:5 (d5) TG (4.03 µM), 14:0-16:1-14:0 (d5) TG (3.99 µM), 15:0-18:1-15:0 (d5) TG (3.97 µM), 16:0-18:0-16:0 (d5) TG (4.05 µM), 17:0-17:1-17:0 (d5) TG (4.14 µM), 19:0-12:0-19:0 (d5) TG (4.01 µM), 20:0-20:1-20:0 (d5) TG (3.81 µM), 20:2-18:3-20:2 (d5) TG (3.96 µM), 20:4-18:2-20:4 (d5) TG (3.90 µM). The 15:0-18:1-15:0 (d5) TG was used for semi-quantification of endogenous TG lipids. Samples were vortexed thoroughly and incubated with 1,000 rpm shaking at room temperature for 30 min, followed by centrifugation at 14,000 rpm at room temperature for 15 min. Supernatants containing lipids were transferred to new tubes. The remaining pellets were re-extracted with 100 µL of MilliQ water and 400 μL of chloroform:methanol (1:2, v:v) containing 0.01% (w/v) butylated hydroxytoluene (BHT) following incubation and centrifugation as described above. The supernatants from the repetitive extractions were collected and pooled, dried by evaporation under vacuum using a GeneVac miVac sample concentrator (SP Scientific, Warminster, PA, USA) and then reconstituted in isopropanol:methanol:chloroform (4:2:1, v:v:v, containing 0.01% BHT) at a final concentration of 4 μL lipid extract per μg protein.

### Sequential functional group derivatization of aminophospholipids and plamalogen-containing lipids

Derivatization of aminophospholipids (i.e., PE and PS) and plasmalogen-containing lipids followed the method previously reported by Ryan & Reid (2016). Prior to derivatization, a 2.5 mM stock solution of triethyamine (TEA) in chloroform was freshly prepared by adding 3.4 μL TEA to 10 mL of chloroform. A 2.5 mM stock solution of S,S’-dimethylthiobutanoylhydroxysuccinimide ester iodide (^13^C_1_-DMBNHS) was freshly prepared by dissolving 4.87 mg ^13^C_1_-DMBNHS in 5 mL of dimethylformamide (DMF). A stock solution of 3.94 mM iodine was freshly prepared by dissolving 10 mg iodine in 10 mL chloroform. A stock solution of 90 mM ammonium bicarbonate was freshly prepared by dissolving 35.6 mg ammonium bicarbonate in 5 mL of HPLC methanol. A solution of 2:1 (v:v) chloroform:methanol containing 266 μM iodine and 2 mM ammonium bicarbonate was prepared by adding 160 μL of 3.94 mM iodine in chloroform to 1.44 mL chloroform, and 53.3 μL of 90 mM ammonium bicarbonate in methanol to 746.7 mL methanol, then combined and placed in an ice bath. Due to the limitations in sample amounts, no replicate derivatization reactions were performed. 4 μL of brain tissue or BDEV lipid extracts were aliquoted to individual wells of a Whatman® Multi-Chem™ 96-well plate (Sigma Aldrich, St. Louis, MO, USA). The solvent was evaporated under vacuum with a GeneVac miVac sample concentrator. 40 μL of a solution of 39:1.1:1 (v:v:v) chloroform:2.5 mM TEA:2.5 mM ^13^C_1_-DMBNHS reagent was added to each dried lipid extract and the 96-well plate was sealed with Teflon Ultra Thin Sealing Tape. Samples were then incubated at room temperature with gentle shaking for 30 min. After incubation, the solvents were evaporated under vacuum with a GeneVac miVac sample concentrator and samples were chilled on ice for 10 min prior to addition of 40 μL of the 2:1 (v:v) chloroform:methanol containing 266 μM iodine and 2 mM ammonium bicarbonate. Reactions were mixed by careful pipetting and the plate was sealed with aluminium foil and then placed on ice for 5 min before solvents were completely removed by evaporation under vacuum with a GeneVac miVac sample concentrator. The dried lipid extracts were washed three times with 40 μL of 10 mM aqueous ammonium. Remaining traces of water were then removed by evaporation under vacuum with a GeneVac miVac sample concentrator. The derivatized brain tissue lipid extracts and BDEV lipid extracts were then resuspended in 50 μL and 25 μL of isopropanol:methanol:chloroform (4:2:1, v:v:v) containing 20 mM ammonium formate respectively. The 96-well plate was then sealed with Teflon Ultra Thin Sealing Tape prior to mass spectrometry analysis.

### Direct infusion nano-electrospray ionization (nESI) - ultrahigh resolution accurate mass spectrometry (UHRAMS) and higher energy collision induced dissociation tandem mass spectrometry (HCD-MS/MS) shotgun lipidome analysis

For underivatized samples, 4 μL of brain tissue or BDEV lipid extracts were aliquoted in triplicate to individual wells of a twin-tec® 96-well plate (Eppendorf, Hamburg, Germany). The brain tissue lipid extracts and BDEV lipid extract were dried and then resuspended in 50 μL (brain tissue) and 25 μL (exosome) of isopropanol:methanol:chloroform (4:2:1, v:v:v) containing 20 mM ammonium formate respectively. The 96-well plate was then sealed with Teflon Ultra Thin Sealing Tape prior to mass spectrometry analysis. 10 μL of each underivatized or derivatized lipid sample was aspirated and introduced via nano-ESI to an Orbitrap Fusion Lumos mass spectrometer (Thermo Fisher Scientific, San Jose, CA, USA) using an Advion Triversa Nanomate (Advion, Ithaca, NY, USA) operating with a spray voltage of 1.1 kV and a gas pressure of 0.3 psi in both positive and negative ionization modes. For MS analysis, the RF lens was set at 10%. Full scan mass spectra were acquired at a mass resolving power of 500,000 (at 200 m/z) across a m/z range of 350 – 1600 using quadrupole isolation, with an automatic gain control (AGC) target of 5e5. The maximum injection time was set at 50 ms. Spectra were acquired and averaged for 3 min. Following initial ‘sum-composition’ lipid assignments by database analysis (see below), ‘targeted’ higher-energy collision induced dissociation (HCD-MS/MS) product ion spectra were acquired on selected precursor ions at a mass resolving power of 120,000 and default activation times in positive ionization mode using the underivatized lipid extracts to confirm the identities of lipid head groups, or in negative ionization mode using underivatized lipid extracts for fatty acid chain identification. HCD-MS/MS collision energies were individually optimized for each lipid class of interest using commercially available lipid standards whenever possible.

### Lipid identification, quantification and data analysis

‘Sum composition’ level lipid identifications were achieved using a developmental version of LipidSearch software 5.0α (Mitsui Knowledge Industry, Tokyo, Japan) by automated peak peaking and searching against a user-defined custom database of lipid species (including the deuterated internal standard lipid species and allowing for the mass shifts introduced by ^13^C_1_-DMBNHS and iodine/methanol derivatization). The parent tolerance was set at 3.0 ppm, a parent ion intensity threshold three times that of the experimentally observed instrument noise intensity, and a max isotope number of 1 (i.e., matching based on the monoisotopic ion and the M+1 isotope), a correlation threshold (%) of 0.3 and an isotope threshold (%) of 0.1. The lipid nomenclature used here follows that defined by the LIPID MAPS consortium (Fahy et al., 2005). Semi-quantification of the abundances of identified lipid species was performed using an in-house R script, by comparing the identified lipid ion peak areas to the peak areas of the internal standard for each lipid class or subclass, followed by normalization against the total protein amount in the samples.

### Lipidomic statistical analysis

Data filtering was implemented based on the 4 comparison groups including NC tissue, AD tissue, NC BDEV and AD BDEV. For lipid identifications obtained from underivatized samples, the mean normalized abundances for each lipid species were included only if the lipid was identified in all three measurement replicates, and detected in at least 6 out of 8 biological samples in each group. For lipid identifications obtained from derivatized samples, the mean normalized abundances for each lipid species were included only if the lipid was identified in at least 6 out of 8 biological samples in each group. Differences in the mean normalized abundances, at the lipid category, class, subclass or individual ‘sum-composition’ lipid species levels, in NC tissue vs. AD tissue and NC BDEV vs. AD BDEV, were determined by multiple t test followed by Holm-Sidak method corrected for multiple comparison using GraphPad Prism 8.0 software. Statistical significance was set at P < 0.01.

## Results

### Characterization of BDEVs

We recently reported a method to isolate small extracellular vesicles from post-mortem human frontal cortex, that contain the hallmarks of endosomal derived exosomes (Cheng et al., 2020, Vella et al., 2017). These vesicles fulfilled the experimental requirements as set out by the International Society for Extracellular Vesicles 2018 guidelines (Théry et al., 2018). Here, we employed this method to isolate BDEVs from post-mortem frontal cortex brain tissue obtained from AD and NC subjects. Dissociated collagenase treated tissue (“Brain+C”) was subject to sequential centrifugation and the 10,000 x g supernatant loaded on top of a triple sucrose density fraction to separate EVs based on density (**Supplemental Figure 1**). Factors such as post-mortem delay and storage can negatively impact tissue quality, resulting in contamination of EV pellets with cellular debris (Vella et al., 2017). We previously reported that immunoblotting provides the most robust quality-control measure for tissue EV isolation, with proteins such as calnexin providing a useful indicator of EV purity. Consequently, all BDEVs isolated as part of the current study were subject to western blotting and TEM (representative images in **Figures 1A** and **1B**) to screen for EV markers and contaminants before subjecting samples to downstream lipidomic analysis. As per our previous studies (Vella et al., 2017), small EVs were identified in fraction 2 (F2) with a density of approximately 1.08 g/cm^3^. EVs in F2 were enriched in the small EV-specific markers TSG101 and syntenin, and depleted of calnexin (**Figure 1A**). TEM images of F2 showed small, cup-shaped membrane vesicles with a diameter of 40-200 nm for both AD and NC samples (**Figure 1B**). Together, these results suggest that F2 contains vesicles that are consistent with the density, morphology, size and protein co-enrichment of endosome derived small BDEVs as we have previously reported. **Supplemental Table 2** shows the protein content determined for each tissue and BDEV fraction, where on average, AD subjects yielded approximately half the BDEV yield compared to NC subjects, a possible reflection of the cellular dysfunction occurring in late stage disease.

**Figure 1.**
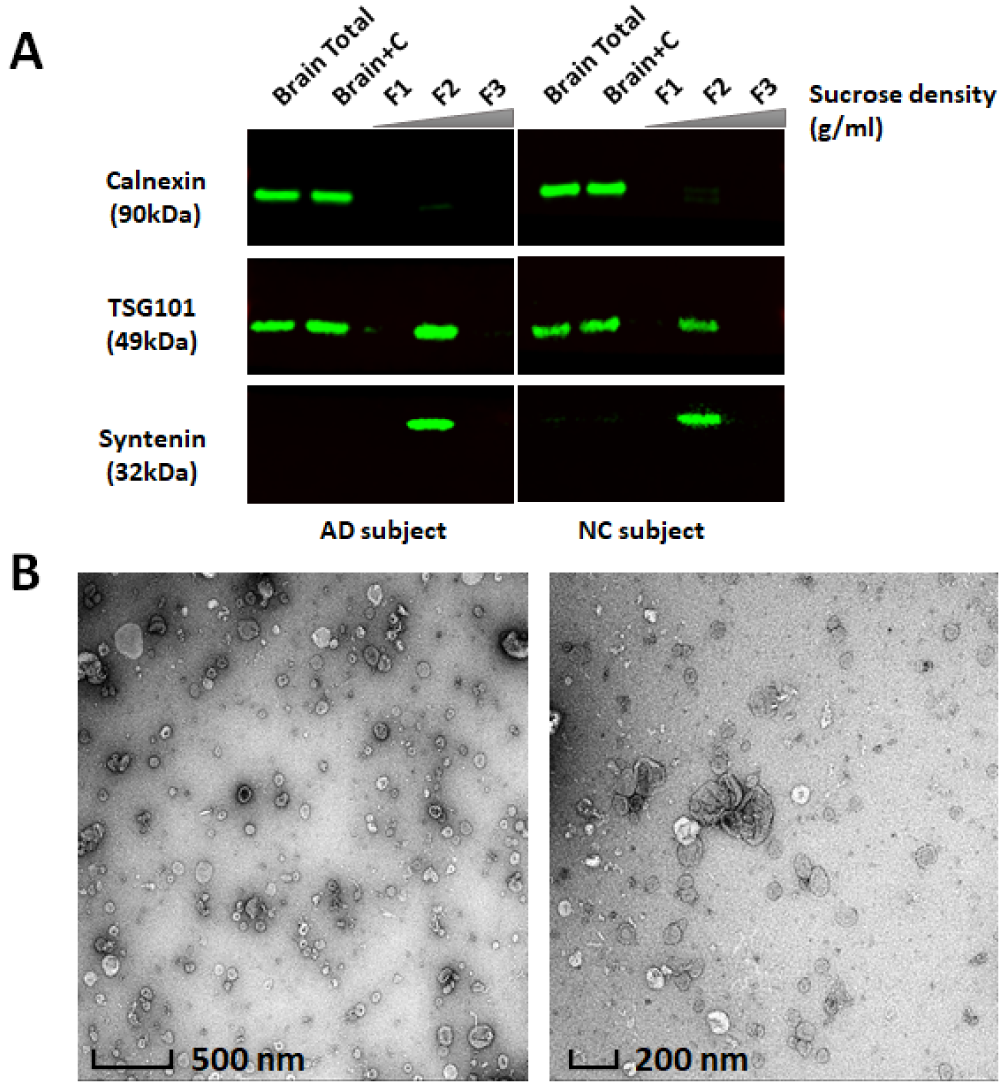
Characterization of BDEVs from AD or NC frontal cortex. **(A) Western blot analysis.** Equivalent amount of protein from human frontal cortex brain homogenates (Brain Total), brain homogenates after collagenase treatment (Brain+C) and BDEV suspensions (F1, F2, and F3) were subjected to SDS-PAGE. Total proteins were visualized using stain-free technology to ensure similar loading. Frontal cortex brain tissue homogenates, ‘Brain Total’ and ‘Brain+C’, were enriched in calnexin, while calnexin was not detectable in an equivalent amount of BDEV protein. Proteins typical of endosome derived exosomes, TSG101 and syntenin, were observed in F2, illustrating that F2 is enriched in exosome-like vesicles. The densities in F1, F2 and F3 were approximately 1.02 g/ml, 1.08 g/ml and 1.17 g/ml respectively. Immunoblots images are representative of 8 independent NC and 8 AD human tissue samples. **(B) Transmission electron microscopy (TEM) of NC BDEV**. All BDEVs from F2 were fixed with 1% (w/v) glutaraldehyde, negatively stained with 2% (w/v) uranyl acetate and visualized by a FEI Tecnai F30 transmission electron microscope. The zoomed-out image (left) provides an overview of the BDEV suspensions with scale bar representing 500 nm. The close-up image (right) shows clearer small, cup-shaped BDEVs which is consistent with the morphology of exosomes with scale bar representing 200 nm. The TEM images of BDEVs in F2 are representative of the BDEVs from all samples (AD and NC). AD, Alzheimer’s disease, NC, neurological control, BDEV, brain derived extracellular vesicles.

### Lipidome analysis of frontal cortex and derived BDEVs in AD

To investigate the lipid composition of frontal cortex tissue and their BDEVs, an in-depth semi-quantitative mass spectrometric analysis was performed on AD and NC tissues, and the BDEVs derived from said tissue. To ensure that downstream analysis was conducted on highly reproducible and confidently identified lipid species, only lipids that met a stringent filtering criteria including identification in all technical replicates and detection in at least 6 out 8 biological samples in each group were included in this analysis. In total, 692 lipid molecules from four main lipid categories, including glycerophospholipids (GP), sphingolipids (SP), glycerolipids (GL) and sterol lipids (ST), covering 17 lipid classes, were identified and semi-quantified at the ‘sum composition’ level (i.e., where the identity of the lipid class, subclass and the total number of carbon atoms and total number of C=C double bonds in the fatty acyl/alkyl/alkenyl moieties are assigned. For example, PE(P-38:4) indicates a 1-(1Z-alkenyl), 2-acylglycerophosphoethanolamine lipid containing a total of 38 carbons and 4 double bonds in the alkenyl and acyl chains). Initial lipidome analysis comparing “Brain Total” and “Brain+C” indicated that collagenase did not result in any significant alterations in the number or abundance of the identified lipids in brain tissue (data not shown). Therefore, all subsequent comparisons of brain tissue and BDEVs were performed using “Brain+C” samples. The number of identified lipid species at the lipid category, class or subclass levels, and their mean summed lipid abundances, either normalized to tissue weight (i.e., pmol/mg tissue), protein content (i.e., pmol/µg protein), total identified lipid concentration (i.e., mol% total lipid), total identified lipid-category concentration (i.e., mol% category), or total identified lipid class concentration (i.e., mol% class) are summarised in **Supplemental Table 3**. A complete list of the individual identified lipids and their abundances from the n=8 AD and NC tissue samples and BDEVs can be found in **Supplemental Table 4**.

Comparison between AD and NC tissue was initially performed using the absolute lipid abundances normalized to tissue (i.e., pmol/mg tissue) (**Supplemental Table 3** and **Supplemental Figure 2**), where no differences were observed at either the total lipid level or at the lipid category or lipid class levels of annotation. However, due to the substantial size difference between cells and BDEVs, it was not possible to perform comparisons between tissue and BDEVs using absolute lipid abundance normalized to tissue, or protein. Therefore, the analysis of differences between tissue and BDEVs was performed by normalizing the concentrations of identified lipids for a given sample to either the total lipid concentration (i.e., mol% total lipid), or total lipid-class concentration (i.e., mol% total lipid class). Furthermore, it was not possible to directly compare lipid abundances between AD and NC BDEVs using absolute lipid abundances normalized to their respective tissue amounts because of the significant difference in BDEV yields (**Supplemental Table 2**), so absolute lipid abundances normalized to protein (i.e., pmol/µg protein), total lipid concentration (i.e., mol% total lipid), and total lipid class concentration (i.e., mol% class) were used.

**Figure 2A** shows the mean of the summed lipid abundances at the lipid category level for AD and NC frontal cortex, and their BDEVs. Features common to BDEVs included significantly higher GP (p <0.0001, mol%) and lower SP (p <0.0001, mol%) compared to parental tissue, independent of diagnosis. At the lipid class level, BDEVs were significantly enriched in PS (p <0.0001, mol%) with a corresponding decrease in ganglioside lipids (p <0.0001, mol%) (**Figure 2B**). Encouragingly, the BDEVs were not abundant in GL or cholesterol ester (CE) lipids, suggesting limited lipoprotein contamination (Serna et al., 2015, Sun et al., 2019, Wang and Eckel, 2014). Cardiolipins (CL), which reside on the inner membrane of mitochondria, was also limited in abundance, further validating the methodological approach we used here to isolate EVs from tissue.

**Figure 2.**
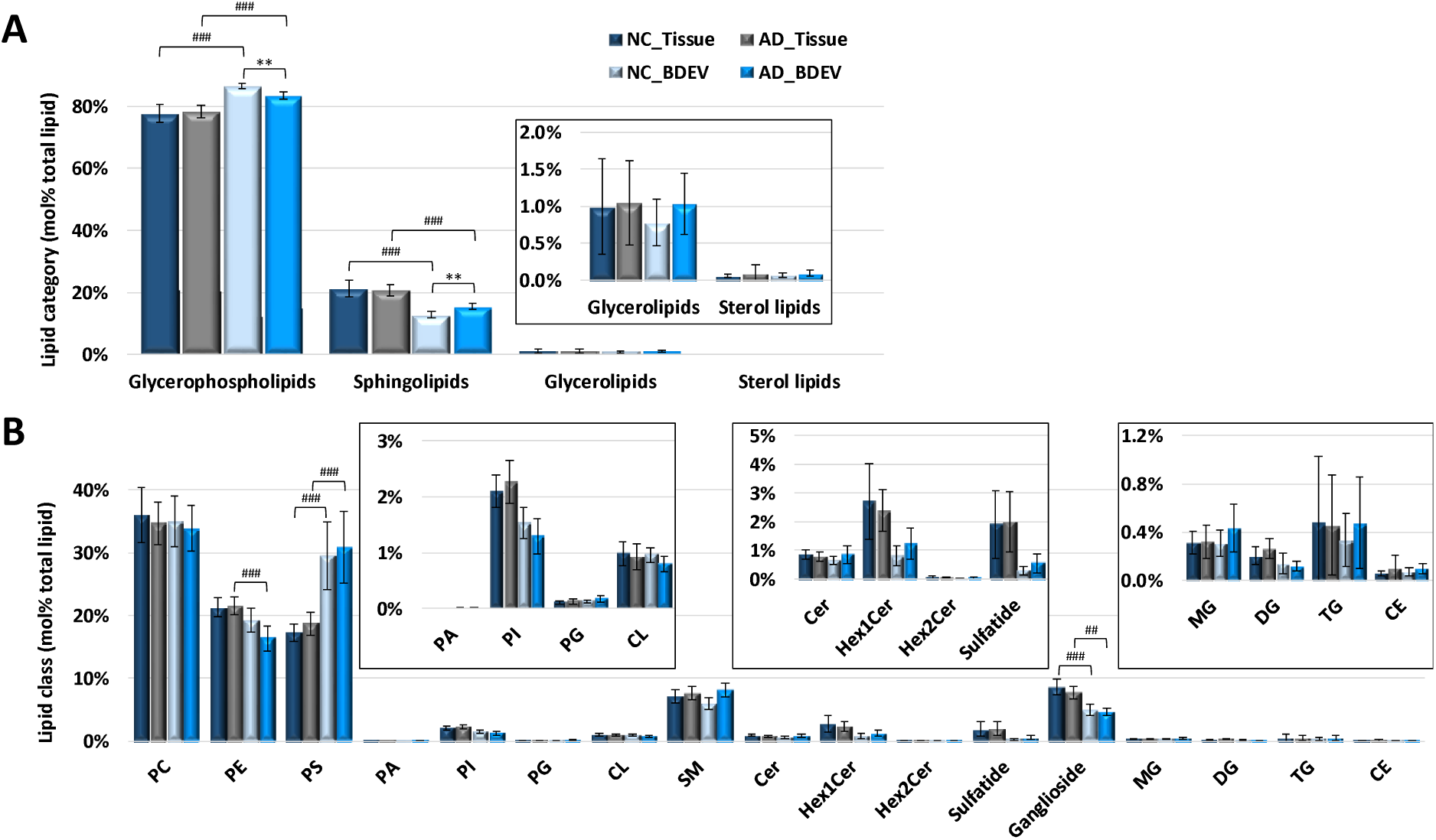
Comparison of mol% total lipid abundance differences between tissue and BDEVs from neurological control (NC) versus Alzheimer’s disease (AD). **(A) Mol% total lipid abundance distributions at the lipid category level.** Four lipid categories, covering glycerophospholipids (GPs), sphingolipids (SPs), glycerolipids (GLs) and sterol lipids (STs) were included in this study. The inset shows the low abundant GL and ST categories for clarity. BDEVs contained significantly higher levels of GPs and corresponding lower levels of SPs in comparison to tissue. No significant difference was found between AD vs. NC tissue while GPs were significantly decreased in AD vs. NC BDEV, with a corresponding significant increase in SPs. **(B) Mol% total lipid abundance distributions at the lipid class level**. A total of 17 lipid classes were identified in this study. The inset shows the low abundant PA, PI, PG, CL, Cer, Hex1Cer, Hex2Cer, sulfatide, MG, DG, TG and CE classes for clarity. BDEVs were found to be significantly enriched in PS lipids, making up approx. 30% of the total lipid abundance, compared to tissue (approx. 17%). Ganglioside lipids were significantly downregulated in BDEV compared to tissue. Data represent the average mol% total lipid abundances ± standard deviation. Statistical significance was determined using multiple t test following correction with Holm-Sidak method, * represents comparisons between AD vs. NC in either tissue or BDEV, # represents comparisons between tissue vs. BDEV in either AD or NC subjects. *p value < 0.01, ** p value < 0.001, *** p value < 0.0001, ^#^ p value < 0.01, ^##^ p value < 0.001, and ^###^ p value < 0.0001. NC, neurological control, AD, Alzheimer’s disease, BDEV, brain derived extracellular vesicles. N = 8 AD subjects and N = 8 NC subjects.

While no differences were observed in lipid category abundance or at the lipid class level in AD frontal cortex tissue relative to NC, presumably due to limited sample size, BDEVs in AD were significantly different (**Figure 2A**). GP was decreased (p < 0.001) with a corresponding increase in SP in AD BDEVs relative to NC BDEVs (p < 0.001) (mol% **Figure 2A** and pmol/ug **Supplementary Table 3**). Due to these significant differences, we investigated the composition of the most abundant lipid classes and subclasses within these categories (**Figure 3**). For the GP category, this included PC, PE and PS lipids at the lipid-subclass (i.e., diacyl, alkylether (O), plamalogen (P), lysoacyl (L), lysoether (L-O) and lysoplasmalogen (L-P)) (**Figures 3A, C** and **E**) and ‘sum-composition’ levels (**Figures 3B, D** and **F**) of annotation, and for the SP category this included SM and Cer lipids at ‘sum-composition’ levels (**Figures 4A** and **4B**, respectively).

**Figure 3.**
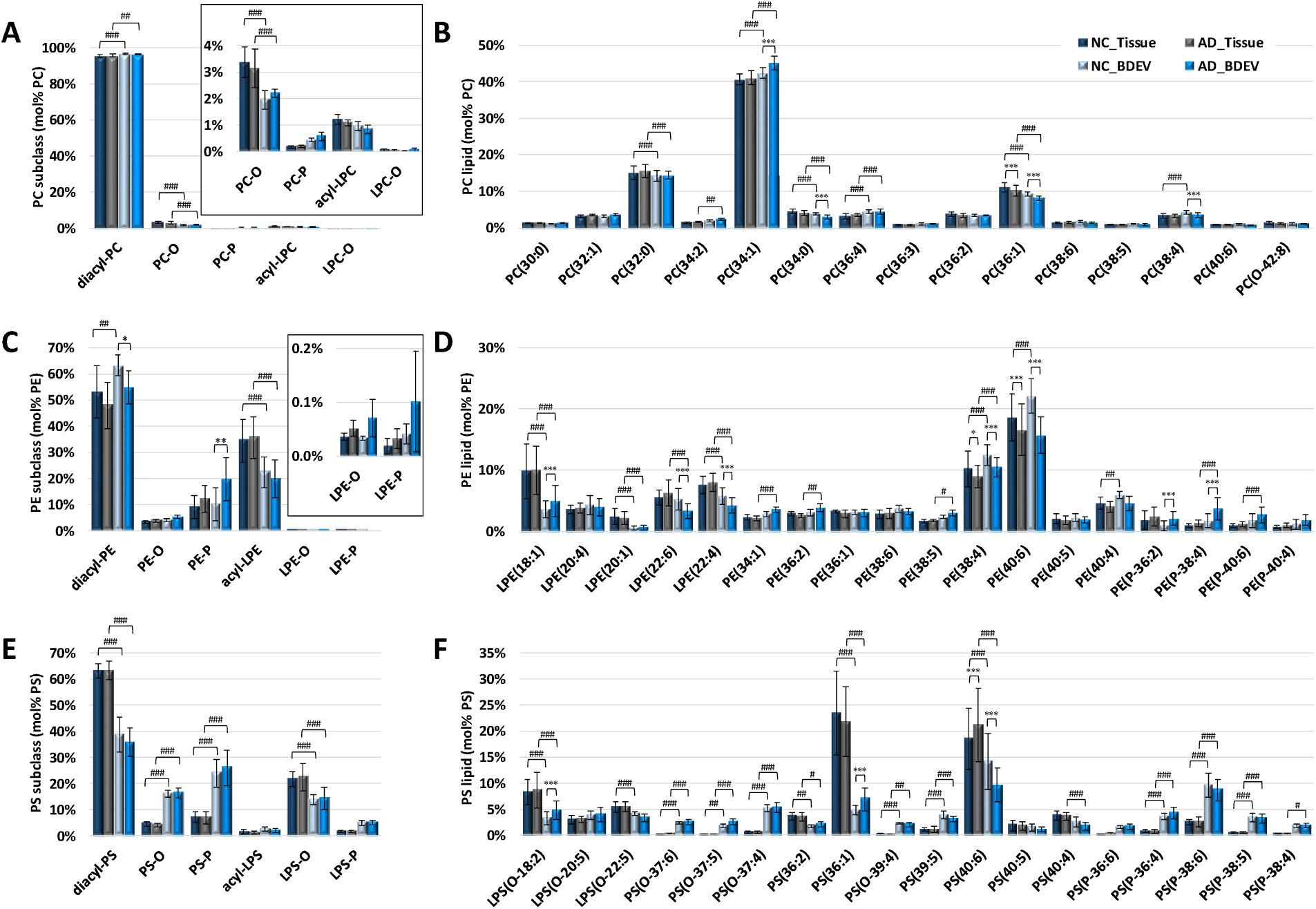
Comparison of PC, PE and PS lipid subclasses abundance (mol% class) and individual lipid molecules (mol% class) between tissue and BDEVs from neurological control (NC) versus Alzheimer’s disease (AD). **(A) Mol% total PC lipid subclass abundance distributions.** The inset shows the low abundant PC-O, PC-P, acyl-LPC and LPC-O for clarity. Significant increase in diacyl-PC was observed in BDEV vs. tissue, accompanied with a decrease in PC-O. No change was observed among PC subclasses between AD vs. NC BDEV. **(B) Mol% total PC lipid abundance distributions of individual PC molecules**. PC(32:0), PC(34:0) and PC(36:1) were observed to be decreased while PC(34:1) and PC(36:4) were observed to be increased in BDEV. Significant increases in PC (32:1) and PC (34:1) and corresponding decrease in PC(34:0), PC(36:1), PC(38:6) and PC(38:4) were observed in AD vs. NC BDEV. **(C) Mol% total PE lipid subclass abundance distributions**. The inset shows the low abundant LPE-O and LPE-P for clarity. Acyl-LPE is decreased in BDEVs relative to tissue. Significant increase in diacyl-PE was observed in BDEV vs. tissue in NC subjects, diacyl-PE was also increased in BDEV vs. tissue in AD, albeit not significant The PE subclass profile revealed differences which distinguish AD BDEVs from NC BDEVs, with diacyl-PE decreased and PE-P increased in AD BDEV. **(D) Mol% total PE lipid abundance distributions of individual PE molecules**. An overall decrease in LPE (LPE(18:1), LPE(20:1), LPE(22:4)) was observed in BDEV relative to tissueLPE(18:1) was increased in AD BDEV realative to NC BDEV. A group of polyunsaturated fatty acid (PUFA) containing PE molecules, including LPE(22:6), LPE(22:4), PE(38:4) and PE(40:6), were found to be significantly decreased in AD BDEV. A group of the most abundant PE-P lipids, including PE(P-36:2) and PE(P-38:4), was also significantly increased in AD BDEV. **(E) Mol% total PS lipid subclass abundance distributions**. A significant decrease was observed in diacyl-PS and LPS-O in BDEV relative to tissue, accompanied with an enrichment of PS-O and PS-P. **(F) Mol% total PS lipid abundance distributions of individual PS molecules**. An overall increase in ether PS species, including PS(O-37:6), PS(O-37:5), PS(O-37:4), PS(O-39:4), PS(P-36:4), PS(P-38:6), PS(P-38:5) and PS(P-38:4), was observed in BDEV relative to tissue. LPS(O-18:2) and PS(36:1) were significantly upregulated in AD vs. NC BDEV while PS(40:6) was significantly decreased. PS did not show distinct difference in ether species between AD and NC. Only the most abundant lipid molecules in each lipid class are shown for clarity. Data represent the average mol% total lipid class abundance ± standard deviation. Statistical significance was determined using multiple t test following correction with Holm-Sidak method, * represents comparisons between AD vs. NC in either tissue or BDEV, # represents comparisons between tissue vs. BDEV in either AD or NC subjects. *p value < 0.01, ** p value < 0.001, *** p value < 0.0001, ^#^ p value < 0.01, ^##^ p value < 0.001, and ^###^ p value < 0.0001. NC, neurological control, AD, Alzheimer’s disease, BDEV, brain derived extracellular vesicles. N = 8 AD subjects and N = 8 NC subjects.

**Figure 4.**
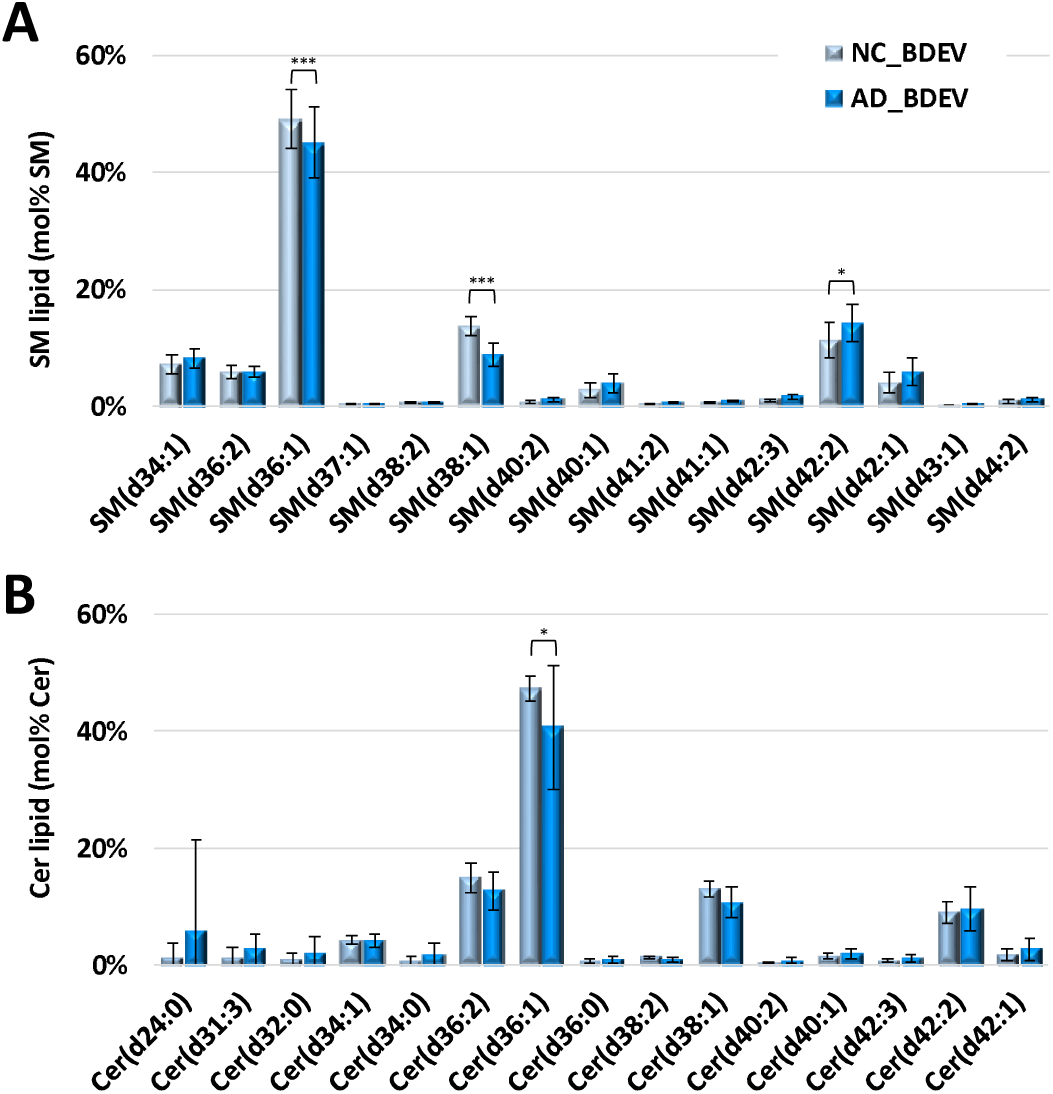
Comparison of SM and Cer individual lipid molecules (mol% class) of BDEV from neurological control (NC) versus Alzheimer’s disease (AD) (n=8 each). **(A) Mol% total SM lipid class abundance distributions.** Significant decreases in SM(d36:1) and SM(d38:1), accompanied with an increase in SM(d42:2), predominantly the SM(d18:1_24:1) species, were observed in AD vs. NC BDEV. **(B) Mol% total Cer lipid class abundance distributions**. Cer(d36:1) was found significantly lower in AD vs. NC BDEV. Only the most abundant lipid molecules in each lipid class are shown for clarity. Data represent the average mol% total lipid class abundance ± standard deviation. Statistical significance was determined using multiple t test following correction with Holm-Sidak method, *p value < 0.01, ** p value < 0.001 and *** p value < 0.0001. NC, neurological control, AD, Alzheimer’s disease, BDEV, brain derived extracellular vesicles.

At the GP-subclass level of annotation, common features of BDEVs were apparent including significant increases in mol% diacyl-PC (**Figure 3A**), diacyl-PE (**Figure 3C**), and PS-O and PS-P (**Figure 3E**) lipids, and corresponding decreases in mol% PC-O, acyl-LPE, diacyl-PS and LPS-O species relative to tissue. At the mol% ‘sum-composition’ level of annotation BDEVs contained significant increases in PC(34:1) and PC(36:4) lipids, and corresponding decreases in PC(32:0), PC(34:0) and PC(36:1) species (**Figure 3B**), along with more extensive acyl chain remodelling of PE and PS lipids, including increased PE(38:4), PS(O-37:6), PS(O-37:5), PS(O-37:4), PS(O-39:4), PS(39:5), PS(P-36:4), PS(P-38:6), PS(P-38:5) and PS(P-38:4) and decreased LPE(18:1), LPE(20:1), LPE(22:4), LPS(O-18:2), PS(36:1) and PS(40:6) relative to tissue (**Figure 3D** and **F**). The observed increase in certain plasmalogen containing PE and PS lipids is consistent with the results observed from the mol% lipid subclass level of analysis that indicated dramatic remodelling of the BDEV composition to incorporate more alkyl-and alkenyl-containing species (particularly those containing the PS head group) rather than the more typical diacyl-containing species (**Figure 3A, 3C and 3E**).

Comparing NC with AD at the subclass level revealed an approximately two-fold increase in PE-P (mol%) in AD BDEVs relative to NC (**Figure 3C**). At the mol% ‘sum-composition’ level, remodelling of the acyl chain compositions was observed in the frontal cortex, resulting in several lipid species differentiating AD and NC e.g., decreased PE(38:4) and PE(40:6), and increased PS(40:6) (**Figures 3D** and **3F**).

Most notably, significant acyl chain remodelling was observed in AD BDEVs, enabling differentiation from NC samples (mol% ‘sum-composition’ level). These included extensive and significant acyl chain remodelling of PE and PS lipids including increased LPE(18:1), PE(P-34:1), PE(P-36:2), PE(P-38:4), LPS(O-18:2) and PS(36:1), and decreased LPE(22:6), LPE(22:4), PE(38:4), PE(40:6) and PS(40:6) (**Figures 3D** and **3F**), along with small but significant increases in PC(34:1) species, with corresponding decreases in PC(34:0), PC(36:1) and PC(38:4) species (**Figure 3B**). Similar changes in PC, PE and PS lipids were observed when comparing AD and NC BDEVs at the lipid subclass and ‘sum-composition’ levels of annotation, based on absolute i.e pmol/µg protein abundances, with AD BDEVs containing approximately half the amount of LPE(22:6), PE (38:4), PE(40:6) and PE(40:4) relative to NC BDEVs (pmol/ug protein, p<0.0001) (**Supplemental Figure 3**). The observed significant decrease in lipid species in AD BDEVs containing docosahexaenoic acid (C22:6 ω3) e.g., LPE(22:6), PE(40:6) and PS(40:6), docosatetraenoic acid (C22:4 ω6) e.g., LPE(22:4) and arachidonic acid (C20:4 ω6) e.g., PC(38:4) and PE(38:4) (each confirmed by HCD-MS/MS) (**Figures 3D** and **3F**), is consistent with overall remodelling of the lipid compositions to those that contain lower concentrations of polyunsaturated fatty acyl chains.

As seen in **Figure 4**, and in **Supplemental Figures 4** and **5**, remodelling of the composition of several abundant SP lipids also enabled differentiation of AD BDEVs from vs NC BDEVs. For example, comparison of the mol% abundance of individual SM and Cer lipid species in BDEVs from AD vs. NC samples (**Figures 4A** and **4B**, respectively) revealed significant decreases in SM(d36:1), Cer(d36:1) and SM(d38:1) species, and a corresponding increase in the very long chain containing SM(d42:2) (predominantly SM (d18:1/24:1)). Interestingly, this was offset by a significant decrease in the mol% abundance of the sulfatide lipid sulfatide(d42:2) containing the same acyl chain composition (**Supplemental Figure 4C**) in AD BDEVs, suggesting that additional remodelling has occurred within the galactosylceramide pathway of sphingolipid metabolism in AD.

Taken together, these findings demonstrate that AD BDEVs have a unique lipid signature that distinguishes them from BDEVs in the frontal cortex of NC subjects, and highlight the utility of analysing EVs, where significant changes that would otherwise be missed in tissue, are revealed.

## Discussion

Using optimised methods for the enrichment of BDEVs, coupled with deuterated internal standards and direct infusion nano-electrospray ionization (nESI) and ultrahigh resolution accurate mass spectrometry (UHRAMS) analysis (Rustam and Reid, 2018, Wang et al., 2017), this study provides the first semi-quantitative characterisation of the lipid composition of BDEVs derived from CNS tissue. In addition to comprising a lipid signature similar to EVs sourced from other biological materials, frontal cortex-derived EVs were enriched in ether-containing PS lipids, a novel finding with implications for EV structure and function. The utility of BDEVs as a sensitive tool for the detection of lipid dyshomeostasis was highlighted, with significant remodelling of the frontal cortex lipidome revealed in BDEVs in AD in the absence of detectable changes in the parent tissue.

The lipid profiles of EVs have been reported for several cell lines and parasites, and from biological fluids including urine, plasma and serum (Brouwers et al., 2013, Haraszti et al., 2016, Laulagnier et al., 2004, Llorente et al., 2013, Lydic et al., 2015, Simbari et al., 2016, Skotland et al., 2017a, Skotland et al., 2020, Sun et al., 2019). In agreement with these studies (Haraszti et al., 2016, Laulagnier et al., 2004, Llorente et al., 2013, Skotland et al., 2017a, Skotland et al., 2017b, Trajkovic et al., 2008, Chan et al., 2012, Chen et al., 2019, Lydic et al., 2015), we show that human frontal cortex derived EVs are enriched in PS lipids. PS lipids make up a substantial proportion of the EV membrane and are proposed to play a role in facilitating EV uptake by recipient cells (Kastelowitz and Yin, 2014, Laulagnier et al., 2004, Record et al., 2014, Record et al., 2018, Wei et al., 2016, Matsumura et al., 2019, Sharma et al., 2017). Decreased diacyl-PC and higher levels of lysophospholipids, SM, Cer and ganglioside have previously been reported in EVs from urine, cell lines and blood (Haraszti et al., 2016, Llorente et al., 2013, Skotland et al., 2017a, Skotland et al., 2020, Sun et al., 2019, Lydic et al., 2015). These findings were not observed here, likely due to the differences in cell and tissue type between studies.

While PC and PE ethers have been identified previously in EVs, here we reveal that EVs are abundant in alkyl- and alkenyl-(i.e., plasmalogen) ether-containing PS species, supporting the suggestion that ether lipids are a feature of EVs (Llorente et al., 2013, Lydic et al., 2015, Simbari et al., 2016, Skotland et al., 2017a, Skotland et al., 2019). Ether lipids can modulate membrane rigidity and permeability (Dorninger et al., 2017, Glaser and Gross, 1994, Wallner and Schmitz, 2011), possibly mediating EV interaction with host cells and contributing to stability in the environment.

Lipid remodelling, specifically alterations in GP and SP, are associated with AD pathogenesis (Bennett et al., 2013, Han et al., 2011, He et al., 2010, Kosicek and Hecimovic, 2013, Mielke et al., 2010a, Mielke et al., 2010b, Mielke et al., 2011, Mielke et al., 2014, Mielke and Lyketsos, 2010, Wong et al., 2017, Wood, 2012). Down regulation of the glycerophospholipids PC, PE and glycerophosphatidylinositol (PI) in AD, with a general decline in plasmalogen lipids, mainly PC-P and PE-P, has been observed in multiple brain regions in AD (Ginsberg et al., 1995, Grimm et al., 2011, Han et al., 2001, Igarashi et al., 2011, Kosicek and Hecimovic, 2013, Wood, 2012). No significant changes were detected in plasmalogen levels in AD frontal cortex (relative to NC) in this study, however clear differences in ether-PE lipids, including both PE-O and PE-P, were observed in the BDEVs isolated from these tissues. Specifically, plasmalogen PE molecules PE(P-36:2) and PE(P-38:4), were significantly increased in BDEVs in AD. Plasmalogens are abundant in the CNS and are considered an antioxidant due to the presence of a carbon-carbon double bond at the ether linkage that is prone to oxidation (Dorninger et al., 2017, Su et al., 2019, Wallner and Schmitz, 2011). Plasmalogens could play an active role in scavenging oxidative stress and reducing inflammatory responses in recipient cells with studies showing that PE-P can rescue neuronal cell death (Hossain et al., 2013, Wood et al., 2010). Whether donor cells become more susceptible to oxidation following release of PE-P via BDEVs is not yet known, with further studies required to determine their role in disease.

The CNS is highly enriched with poly unsaturated fatty acid (PUFA), with the majority of the PUFA content encapsulated in GP (Bazinet and Laye, 2014, Dyall, 2015). The BDEVs in this study recapitulated this brain-specific feature. Moreover, the C22:6 fatty acyl chain-containing species, LPE(22:6), PE(40:6) and PS(40:6) were found to be decreased ∼two fold in AD BDEVs. The C22:6 is the most abundant PUFA chain in GP and it is derived from docosahexaenoic acid, DHA, which is the precursor of a group of beneficial bioactive anti-inflammatory pro-resolving mediators (Serhan and Levy, 2018, Whittington et al., 2017). Notably, the DHA-derived specialised pro-resolving mediators (SPM) are capable of attenuating Aß amyloidogenesis and enhancing Aß phagocytosis in AD (Fiala et al., 2015, Lukiw et al., 2005). The decreased C22:6 content in AD BDEVs in this study supports DHA deficiency in AD, a known feature of the disease (Morris et al., 2003, Tully et al., 2003) and suggest that’s that the level of C22:6 in BDEVs could serve as a peripheral marker of AD.

As sphingolipid metabolism has also been reported to be remodelled in AD (Haughey et al., 2010, He et al., 2010, Mielke and Lyketsos, 2010, Wood, 2012), we investigated lipid profile changes in SM and Cer lipids. SM is predominantly found on the plasma membrane and is one of the main components of lipid-rafts. The abundances of SM lipids appear to be modified in AD, however there is some contention as to whether SM is decreased or increased, with reports suggesting variability between brain regions (Bandaru et al., 2009, Chan et al., 2012, Cutler et al., 2004, He et al., 2010, Kosicek and Hecimovic, 2013, Pettegrew et al., 2001). SM(d36:1) and SM(d38:1) were decreased in AD BDEVs, accompanied by an increase in very long chain SM (d42:2), predominantly the SM(d18:1_24:1) species.

With the knowledge that lipid metabolism is altered in AD in the CNS, researchers have examined the lipidome of CSF and blood in the search for potential biomarkers, with varied results (Han et al., 2003, Koal et al., 2015, Kosicek et al., 2012, Mielke et al., 2010a, Mielke et al., 2010b, Mielke et al., 2011, Mielke et al., 2014). Anand et al. (2017) discovered that LPC and a group of lipid peroxidation products, including oxidized PC, oxidized-triacylglyceride (TG) and F2-isoprostanes, were up-regulated, while PC, SM, PE, especially PE-P, were found to be declined in AD serum (Gonzalez-Dominguez et al., 2014, Goodenowe et al., 2007, Barupal et al., 2019). PUFA PC species (e.g., PC(16:0/20:5), PC(16:0/22:6), PC(18:0/22:6) etc.) have been reported to be decreased in preclinical AD plasma (Fiandaca et al., 2015, Mapstone et al., 2014, Whiley et al., 2014). The major drawback of these approaches relates to the complexity of the lipidome and measurement of changes not necessarily specific to the CNS or disease which reduces diagnostic accuracy and reproducibility. Other diseases, or co-morbidities such as diabetes and cardiovascular disease, can also contribute to the alteration of the lipid profile, further complicating lipid biomarker discovery.

An ideal lipid peripheral biomarker would ideally be brain-derived to reflect early changes in the CNS in AD. BDEVs can readily cross the blood brain barrier (BBB) (Kanninen et al., 2016, Saeedi et al., 2019, Skoumalová et al., 2011), enabling the study of BDEV lipids in the periphery, effectively eliminating many of the caveats associated with studying complex fluids such as serum or plasma. Our research suggests that future studies should assess ether PE lipids and polyunsaturated fatty acyl containing lipids in peripherally-sourced BDEVs alongside other clinical measures (CSF and neuroimaging assessments) to determine if BDEVs can predict progression from Mild Cognitive Impairment to AD.

## Acknowledgements

This work was supported by grants from the Australian National Health and Medical Research Council (GNT1041413 and GNT1002349 to AFH; 628946 to CLM, KJB, and AFH), the Australian Research Council (LE160100015 to GER), the Bethlehem Griffiths Research Foundation to LJV (Australia), the Alzheimer’s Australia Dementia Research Foundation John Shutes Project Grant to LJV and The Alzheimer’s Association (AARF-18-566256) to LJV (U.S.A). We thank Fairlie Hinton and Geoffrey Pavey from the Victorian Brain Bank.

