## Supplemental Figures and Tables 1 and 2 for "Characterization of Brain-Derived Extracellular Vesicle Lipids in Alzheimer’s Disease"

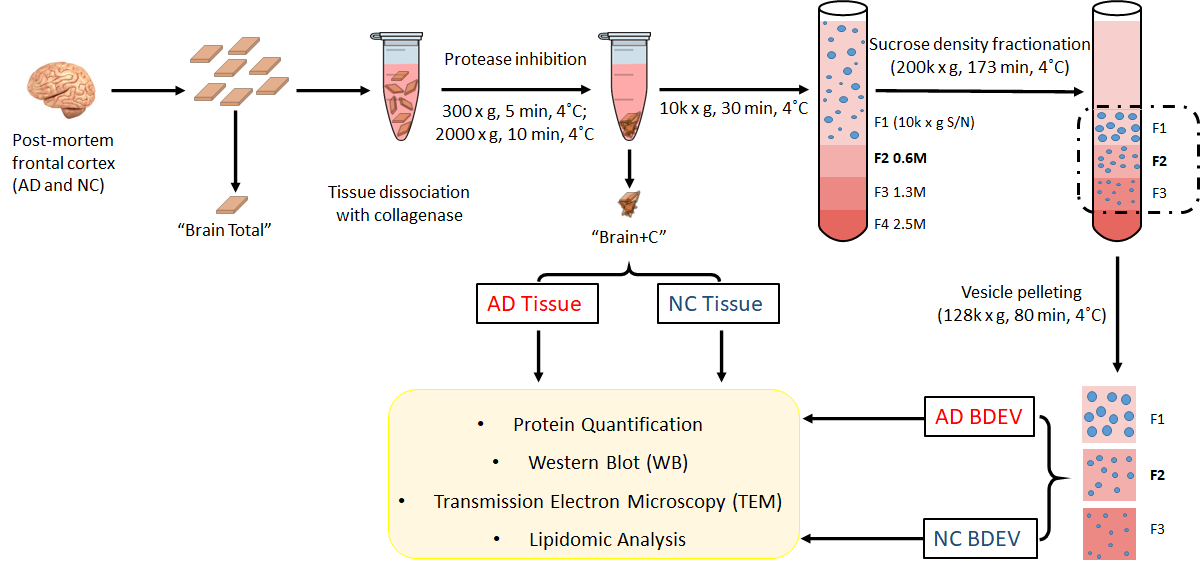

**Supplemental Figure 1. Summary of the experimental workflow for human post-mortem frontal cortex BDEV isolation and analysis.** Fresh frozen frontal cortex AD and NC tissue samples (n=8 each) were sliced on ice. A small section of each was collected as a sample control (“Brain Total”). The remaining sections were dissociated with 50 U/mL of collagenase type 3 in DPBS at 25°C for a total of 20 min with shaking, followed by addition of protease and phosphatase inhibitors and EDTA. The dissociated tissue was spun at 300 x g for 5 min at 4°C (the resulting pellet was referred to as “Brain+C”). The supernatant was transferred to new tube and spun at 2,000 x g for 5 min at 4°C, followed by a 10,000 x g spin for 30 min at 4°C. The extracellular vesicle-containing supernatant was overlaid on a triple sucrose gradient (0.6 M, 1.3 M, 2.5 M) and centrifuged at 200,000 x g (avg) for 173 min at 4°C using a SW40 rotor (Beckman). The vesicles were separated into individual fractions based on density, then each fraction (F1, F2 and F3) was collected and transferred to individual ultracentrifuge tubes then pelleted at 128,000 x g (avg) for 80 min at 4°C using a F37L-8X100 rotor (Thermo Fisher Scientific). “Brain Total”, “Brain+C” and vesicle-enriched fractions, F1, F2 and F3, were subjected to protein analysis (protein quantification and Western blot) and lipidomic analysis. Fraction 2 (F2) was also subjected to size and morphology validation by transmission electron microscopy (TEM). NC, neurological control, AD, Alzheimer’s disease, BDEV, brain derived extracellular vesicles.

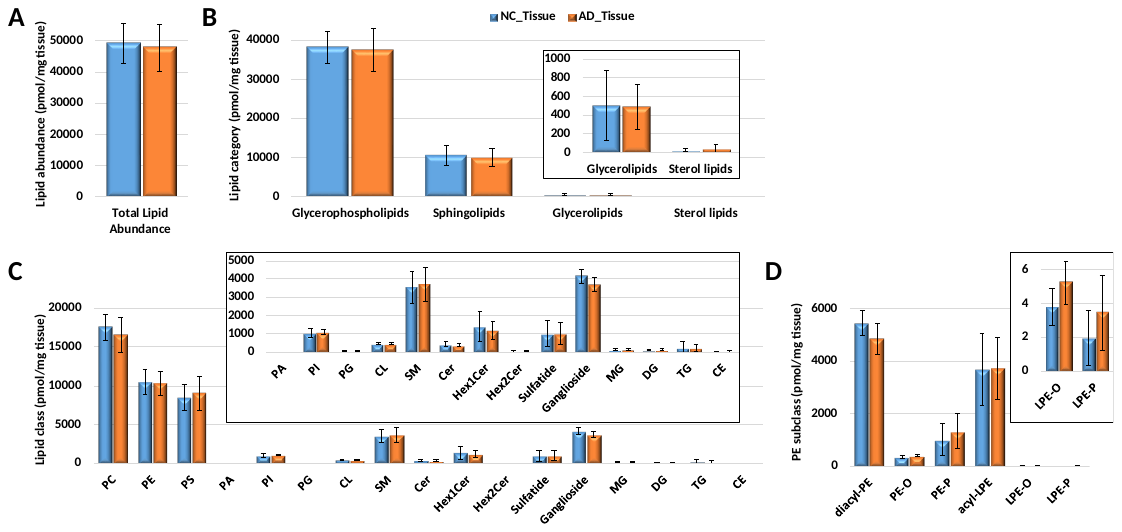

**Supplemental Figure 2. Comparison of lipid abundance (pmol/mg tissue) differences in frontal cortex tissue from neurological control (NC) versus Alzheimer’s disease (AD) (n=8 each). (A) Total lipid abundance**. No clear difference in total lipid abundance between AD vs. NC tissue. **(B) Total lipid abundance at the lipid category level.** Four lipid categories, covering glycerophospholipids (GP), sphingolipids (SP), glycerolipids (GL) and sterol lipids (ST) were included in this study. The inset shows the low abundant GL and ST categories for clarity. **(C) Total lipid abundance (pmol/mg tissue) at the lipid class level.** A total of 17 lipid classes were identified in this study. The inset shows the low abundant PA, PI, PG, CL, SM, Cer, Hex1Cer, Hex2Cer, sulfatide, ganglioside, MG, DG, TG and CE classes for clarity. **(D) Total lipid abundance at the PE subclass level.** The inset shows the low abundant LPE-O and LPE-P for clarity. Data represent the average lipid abundance (pmol/mg tissue) ± standard deviation. Statistical significance was determined using multiple t test following correction with Holm-Sidak method. No significant difference was found. NC, neurological control, AD, Alzheimer’s disease.

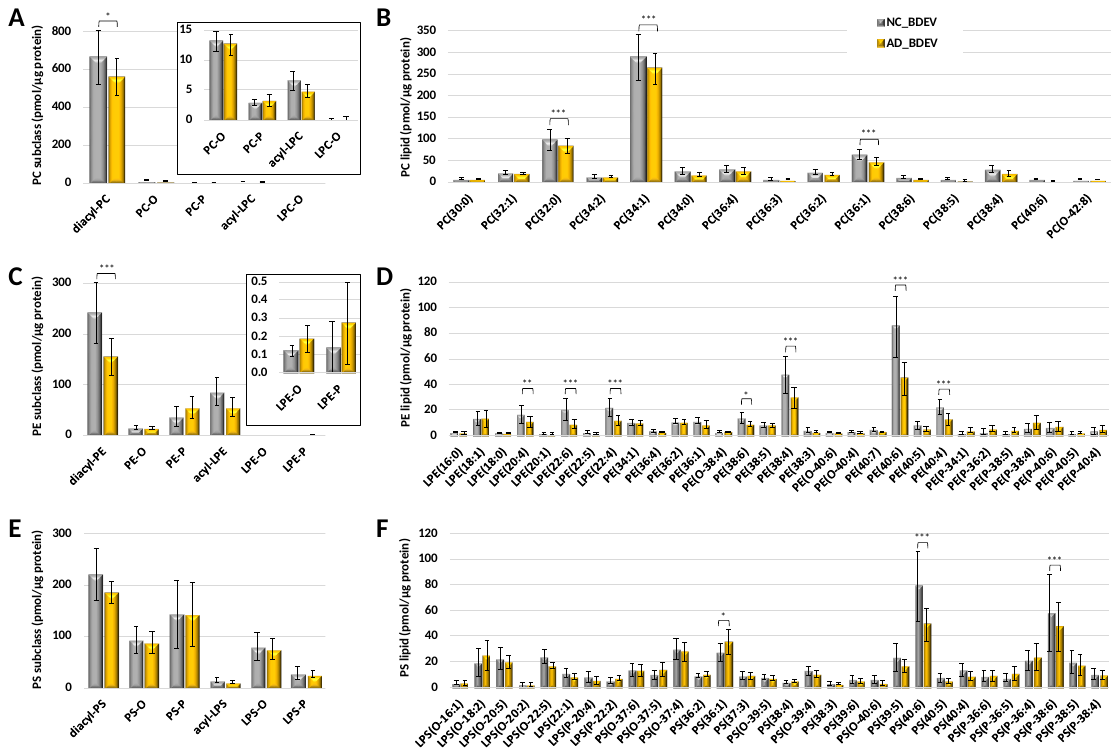

**Supplemental Figure 3. Comparison of PC, PE and PS lipid subclass abundance (pmol/µg protein) and individual lipid molecules (pmol/ µg protein) of BDEV from neurological control (NC) versus Alzheimer’s disease (AD) (n=8 each).** **(A) Total PC subclass abundance.** The inset shows the low abundant PC-O, PC-P, acyl-LPC and LPC-O for clarity. Reduced total diacyl-PC abundance was observed in AD vs. NC BDEV. **(B) Individual PC molecules.** Significant decreases in PC(32:0), PC(34:1), PC(36:1) and PC(38:4) was observed in AD vs. NC BDEV. **(C) Total PE subclass abundance.** The inset shows the low abundant LPE-O and LPE-P for clarity. Reduced total diacyl-PE abundance was observed in AD vs. NC BDEV. Trends toward decreased acyl-LPE and increased PE-P, LPE-O and LPE-P were also observed, albeit not significant. **(D) Individual PE molecules.** A group of polyunsaturated fatty acid (PUFA) containing PE molecules, including LPE(20:4), LPE(22:6), LPE(22:4), PE(38:6), PE(38:4), PE(40:6) and PE(40:4), were found to be significantly decreased and PE(P-38:4) was found to be significantly increased in AD vs. NC BDEV. The signature PE lipid profile could efficiently distinguish AD from NC BDEV, highlighting the potential of using these BDEV as a source of disease biomarkers. **(E) Total PS subclass abundance**. No clear change was observed among PS subclasses to distinguish AD from NC BDEV. **(F) Individual PS molecules.** PS(36:1) was found to be significantly upregulated while LPS(O-22:5), PS(40:6) and PS(P-38:6) were found to be significantly decreased in AD vs. NC BDEV. Only the most abundant lipid molecules in each lipid class are shown for clarity. Data represent the average lipid abundance (pmol/µg protein) ± standard deviation. Statistical significance was determined using multiple t test following correction with Holm-Sidak method, *p value < 0.01, ** p value < 0.001 and *** p value < 0.0001. NC, neurological control, AD, Alzheimer’s disease, BDEV, brain derived extracellular vesicles.

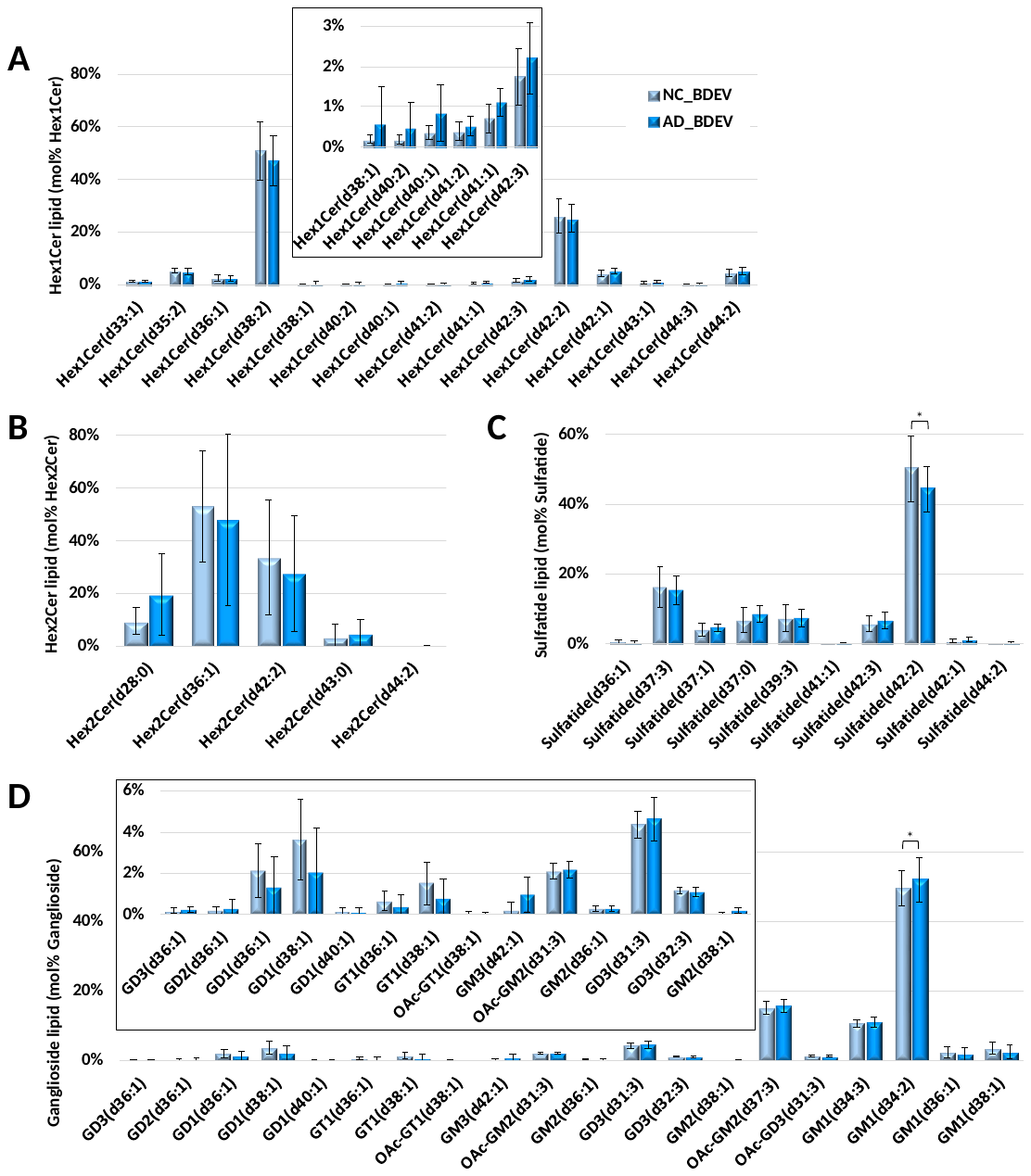

**Supplemental Figure 4. Comparison of Hex1Cer, Hex2Cer, sulfatide and ganglioside individual lipid molecules (mol% class) of BDEV from neurological control (NC) versus Alzheimer’s disease (AD) (n=8 each). (A) Mol% total Hex1Cer lipid class abundance distributions.** **(B) Mol% total Hex2Cer lipid class abundance distributions**. **(C) Mol% total sulfatide lipid class abundance distributions**. Significant decrease in ST(42:2) was observed in AD vs. NC BDEV. **(B) Mol% total ganglioside lipid class abundance distributions**. Only the most abundant lipid molecules are shown for clarity. Data represent the average mol% total lipid class abundance ± standard deviation. Statistical significance was determined using multiple t test following correction with Holm-Sidak method, *p value < 0.01. NC, neurological control, AD, Alzheimer’s disease, BDEV, brain derived extracellular vesicles.

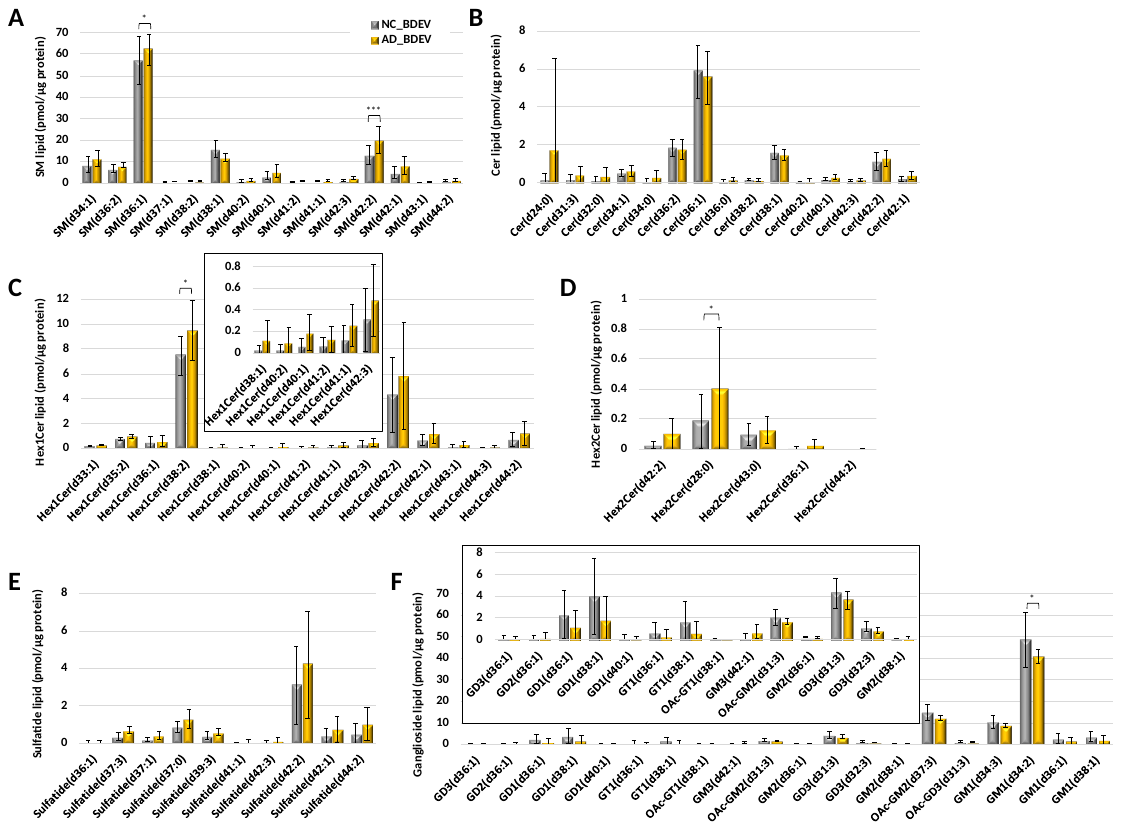

**Supplemental Figure 5. Comparison of individual sphingolipid molecules (pmol/ µg protein) of BDEV from neurological control (NC) versus Alzheimer’s disease (AD) (n=8 each).** **(A) Individual SM molecules. (B) Individual Cer molecules. (C) Individual Hex1Cer molecules. (D) Individual Hex2Cer molecules. (E) Individual sulfatide molecules. (F) Individual ganglioside molecules.** Only the most abundant lipid molecules in each lipid class are shown for clarity. Data represent the average lipid abundance (pmol/µg protein) ± standard deviation. Statistical significance was determined using multiple t test following correction with Holm-Sidak method, *p value < 0.01, ** p value < 0.001 and *** p value < 0.0001. NC, neurological control, AD, Alzheimer’s disease, BDEV, brain derived extracellular vesicles.

**Supplemental Table 1. Subject case information**

| **Disease condition** | **Sex** | **Age**  **(yr)** | **Average age**  **(yr)** | **PMI**  **(hr)** | **Average PMI**  **(hr)** | **ApoE**  **genotype** |
| --- | --- | --- | --- | --- | --- | --- |
| AD | Male | 69 | 74.5 ± 7.0 | 13.5 | 23.3 ± 17.4 | E3/E4 |
|  |  | 72.1 |  | 13 |  | E3/E4 |
|  |  | 89.4 |  | 13 |  | E4/E4 |
|  |  | 76.5 |  | 26 |  | E4/E4 |
|  |  | 74.6 |  | 30 |  | E3/E4 |
|  |  | 76.4 |  | 19 |  | E4/E4 |
|  |  | 66.2 |  | 62.5 |  | E3/E3 |
|  |  | 71.4 |  | 9.5 |  | E3/E4 |
| NC | Male | 78.3 | 73.5 ± 5.9 | 46 | 42 ± 16.3 | E3/E3 |
|  |  | 72.6 |  | 42.5 |  | E3/E3 |
|  |  | 75.6 |  | 46 |  | E3/E4 |
|  |  | 72.6 |  | 20.5 |  | E3/E3 |
|  |  | 66.9 |  | 21 |  | E3/E3 |
|  |  | 63.6 |  | 54.5 |  | E3/E3 |
|  |  | 77.5 |  | 69 |  | E3/E3 |
|  |  | 81 |  | 36.5 |  | E3/E4 |

Data represent individual subject information and the average value ± standard deviations (SD) in either AD or NC subjects. AD, Alzheimer’s disease, NC, neurological control, PMI, post-mortem interval.

**Supplemental Table 2. Protein content of Fraction 2 (BDEV) in individual subject tissue samples.**

| **Disease condition** | **Fresh frozen tissue weight (mg)** | **Tissue protein (µg/mg tissue)** | **Average tissue protein (µg/mg tissue) ^a^** | **Total protein in Fraction 2 (BDEV, µg)** | **Vesicle yield ^b^** | **Average vesicle yield ^c^** | **Refractive index for Fraction 2 (BDEV) *** |
| --- | --- | --- | --- | --- | --- | --- | --- |
| AD | 2042 | 28.0 | 28.3 ± 5.3 | 294.5 | 0.1442 | 0.0785 ± 0.0342 | 1.3643 |
|  | 2038 | 31.5 |  | 222.1 | 0.1090 |  | 1.3644 |
|  | 2305 | 31.9 |  | 206.6 | 0.0897 |  | 1.3638 |
|  | 977 | 28.8 |  | 68.2 | 0.0698 |  | 1.3634 |
|  | 1219 | 35.7 |  | 76.3 | 0.0626 |  | 1.3652 |
|  | 1481 | 20.3 |  | 66.1 | 0.0446 |  | 1.3634 |
|  | 1131 | 21.1 |  | 52.0 | 0.0460 |  | 1.3641 |
|  | 1099 | 29.1 |  | 68.4 | 0.0623 |  | 1.3645 |
| NC | 2283 | 43.1 | 36.5 ± 6.7 | 552.2 | 0.2419 | 0.1564 ± 0.0577 | 1.3627 |
|  | 1918 | 37.0 |  | 232.8 | 0.1214 |  | 1.3631 |
|  | 1988 | 32.6 |  | 485.5 | 0.2442 |  | 1.3636 |
|  | 2000 | 47.0 |  | 351.2 | 0.1756 |  | 1.3612 |
|  | 2015 | 38.1 |  | 236.9 | 0.1176 |  | 1.3614 |
|  | 1614 | 31.2 |  | 170.5 | 0.1056 |  | 1.3622 |
|  | 1213 | 37.4 |  | 134.2 | 0.1106 |  | 1.3646 |
|  | 994 | 25.6 |  | 133.2 | 0.1340 |  | 1.3632 |

^a^ Statistical difference of tissue protein (µg/mg tissue) between AD and NC group was determined using unpaired student’s *t* test, p value = 0.0171. ^b^ Vesicle yield was determined from the protein content in vesicle fraction (F2, BDEV, in µg) as a function of the tissue weight (in mg) used for BDEV enrichment experiment. ^c^ Statistical difference of vesicle yield between AD and NC group was determined using unpaired student’s *t* test, p value = 0.0054. *The refractive index in Fraction 2 (BDEV) corresponds to the density of approximately 1.08 g/ml (Vella *et al*., 2017). Data represent individual subject information and the average value ± standard deviations (SD) in either AD or NC subjects. BDEV, brain-derived extracellular vesicles, AD, Alzheimer’s disease, NC, neurological control.
